## Supplementary material for "Sonic hedgehog signaling directs patterned cell remodeling during cranial neural tube closure": Figure Supplements

### Figure 2 - Supplement 1

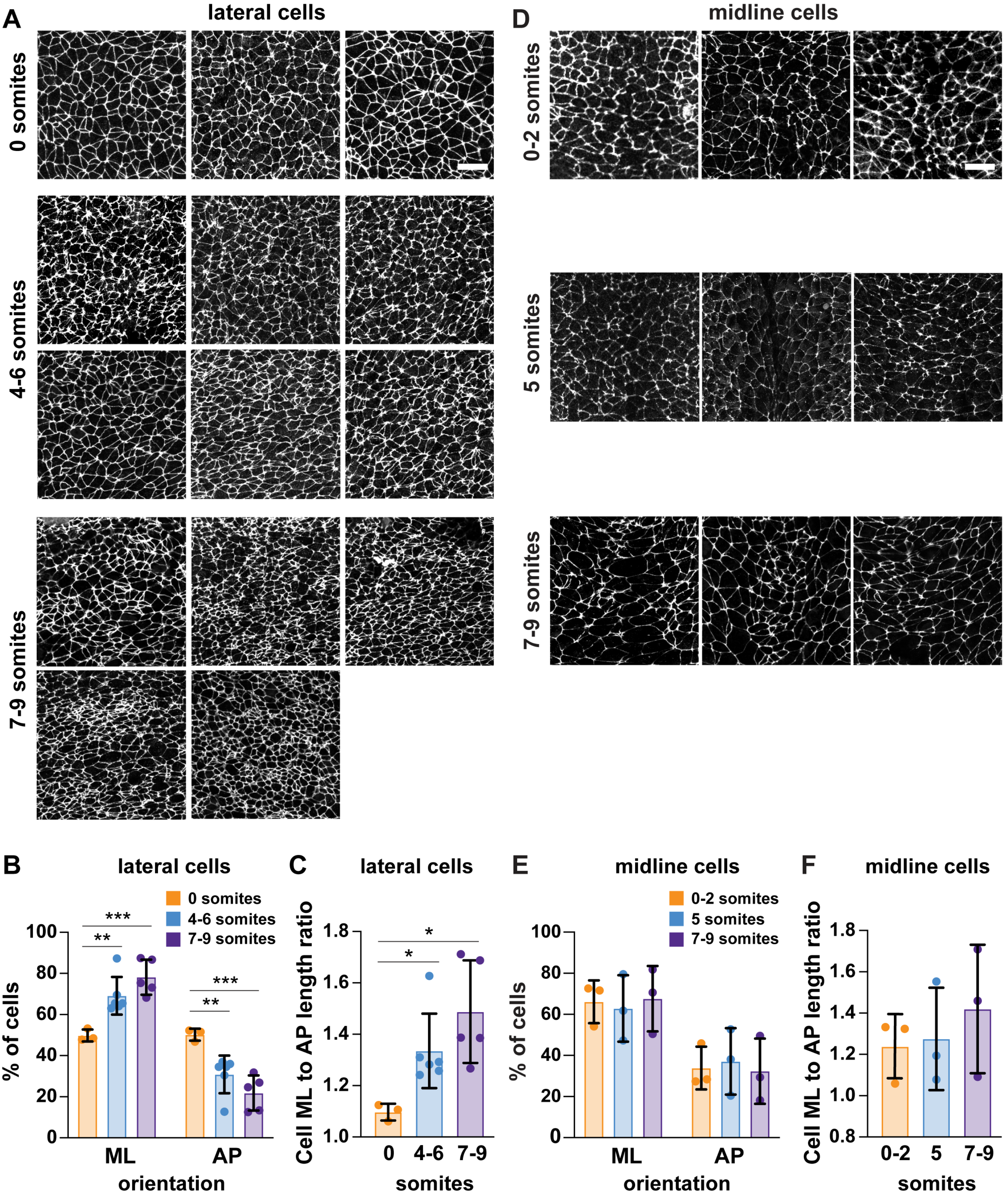

### Figure 3 - Supplement 1

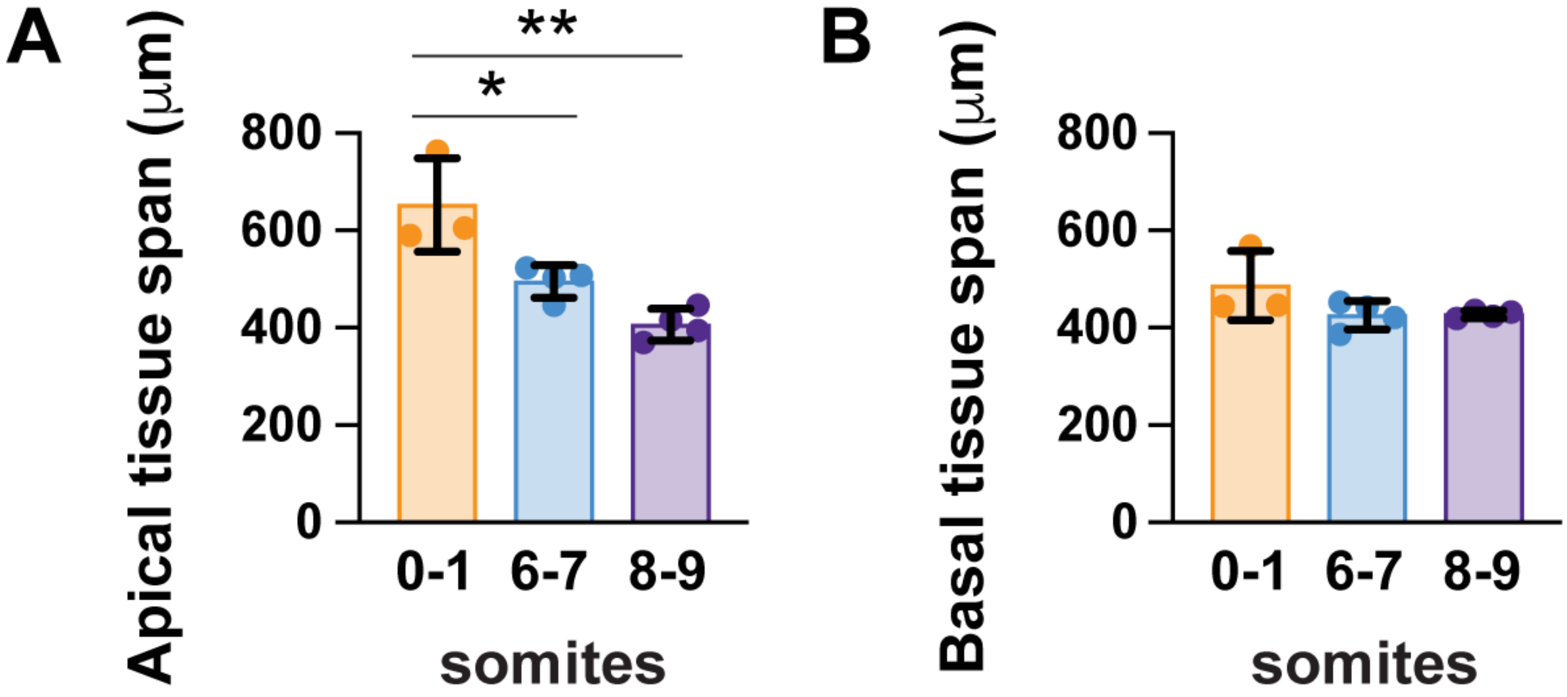

#### Figure 3 - Supplement 2

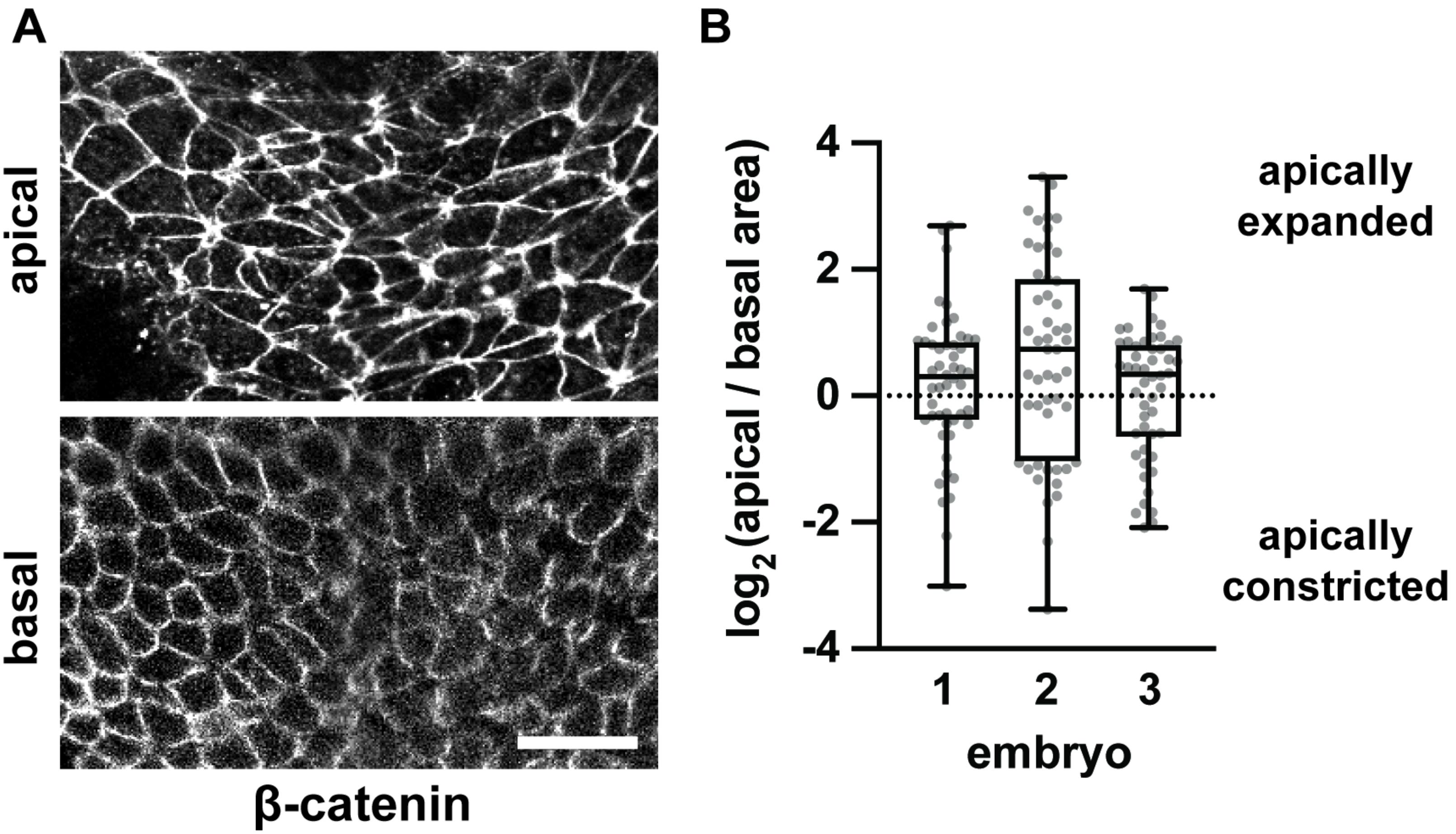

Figure 4 - Supplement 1

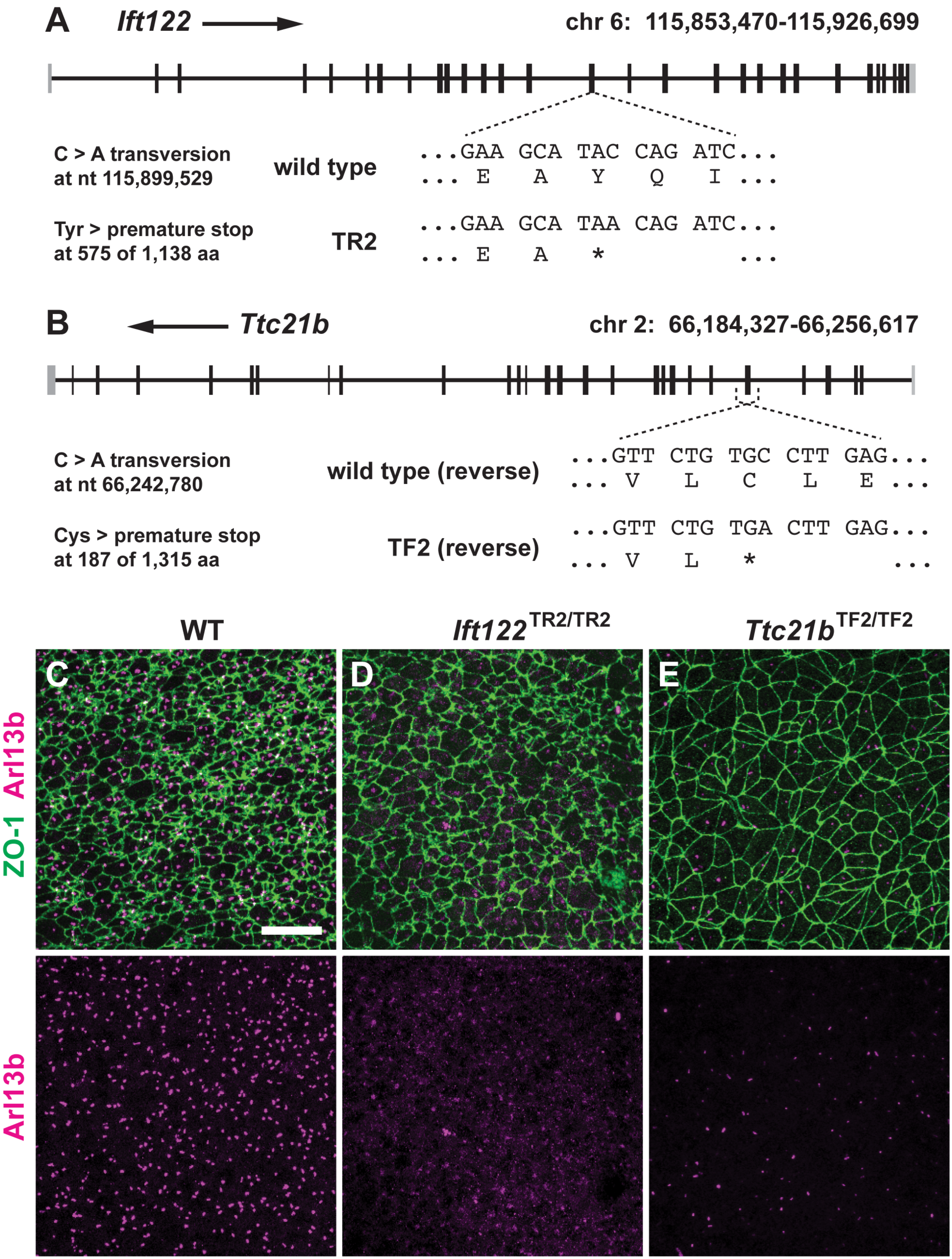

### Figure 4 - Supplement 2

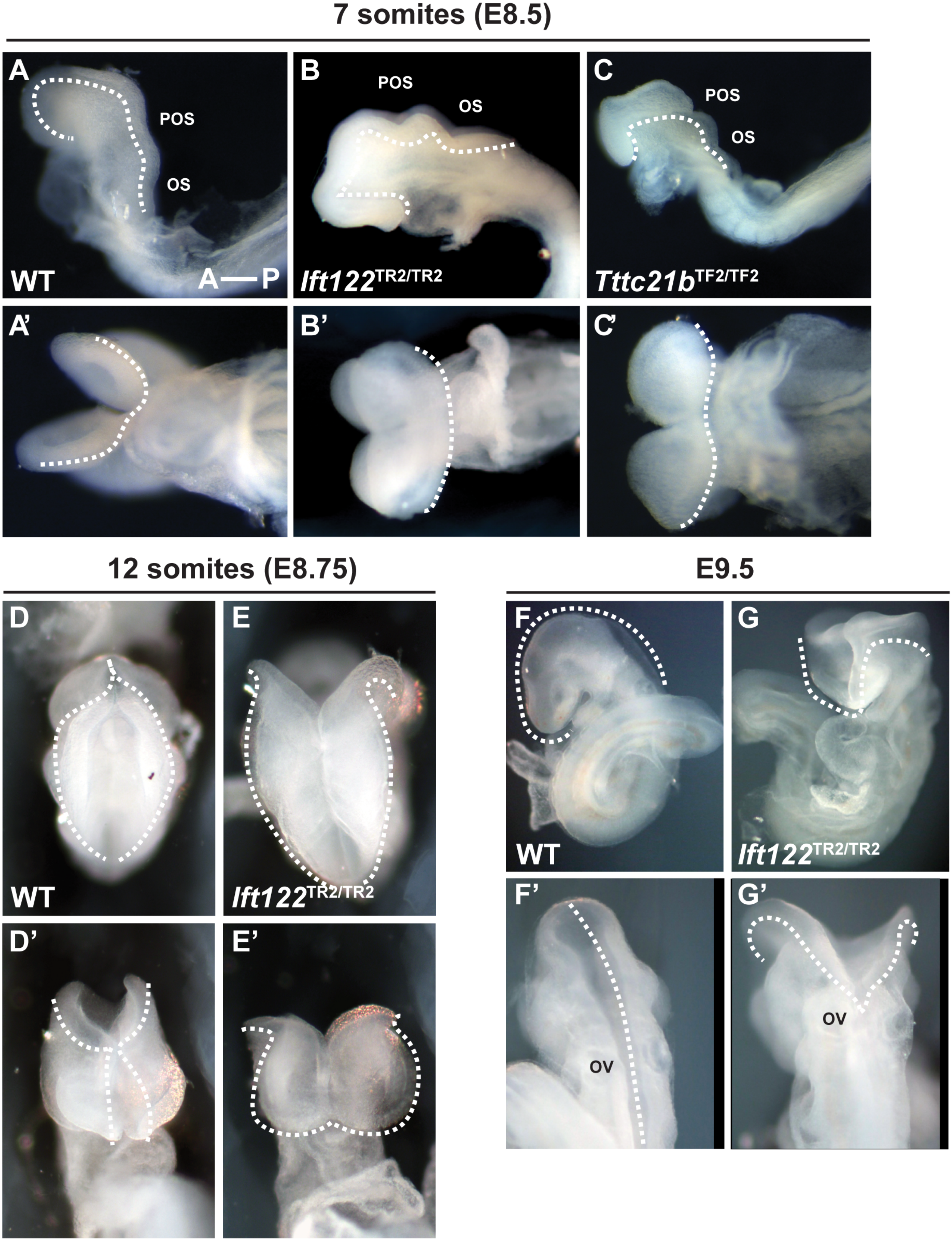

### Figure 4 - Supplement 3

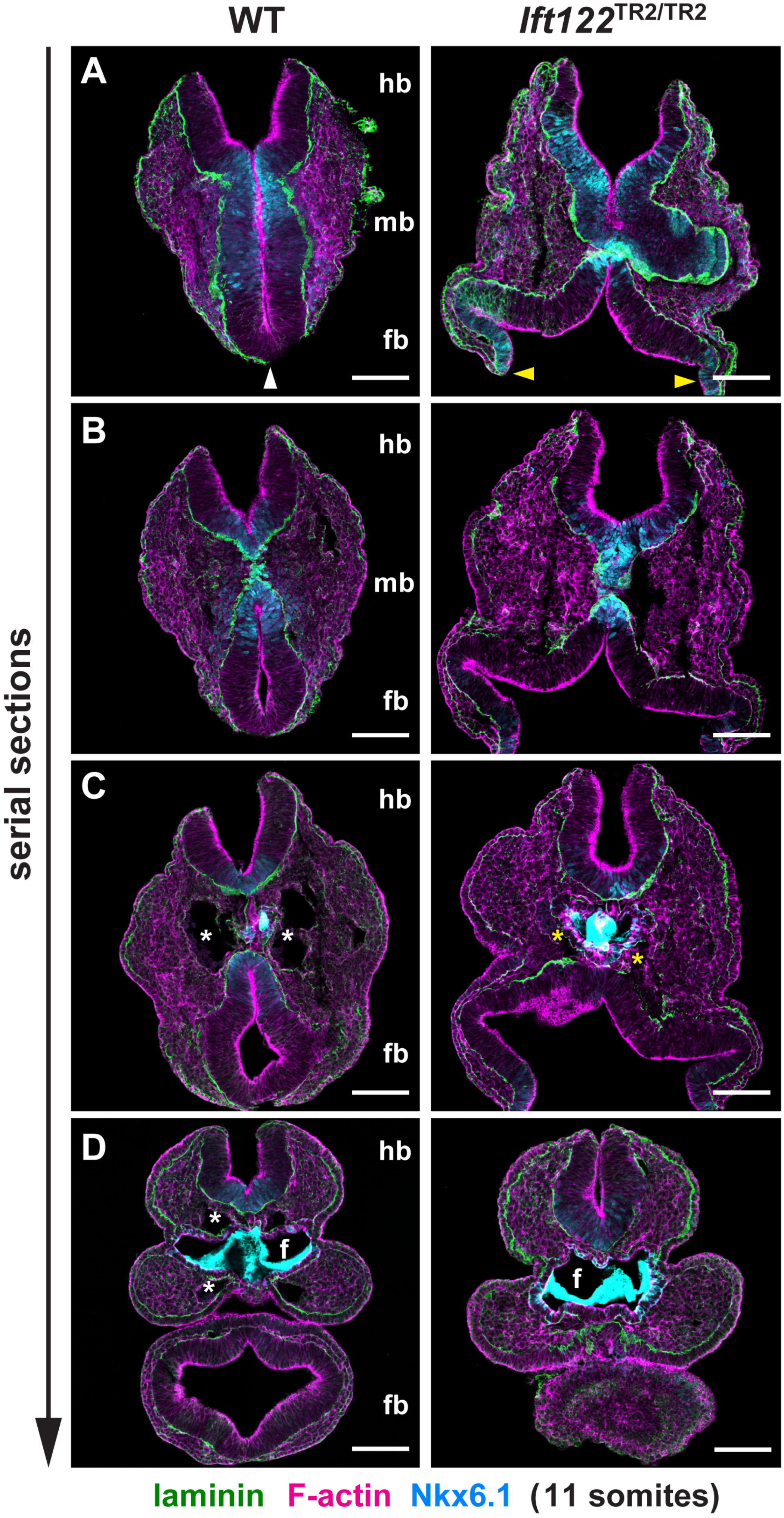

### Figure 4 - Supplement 4

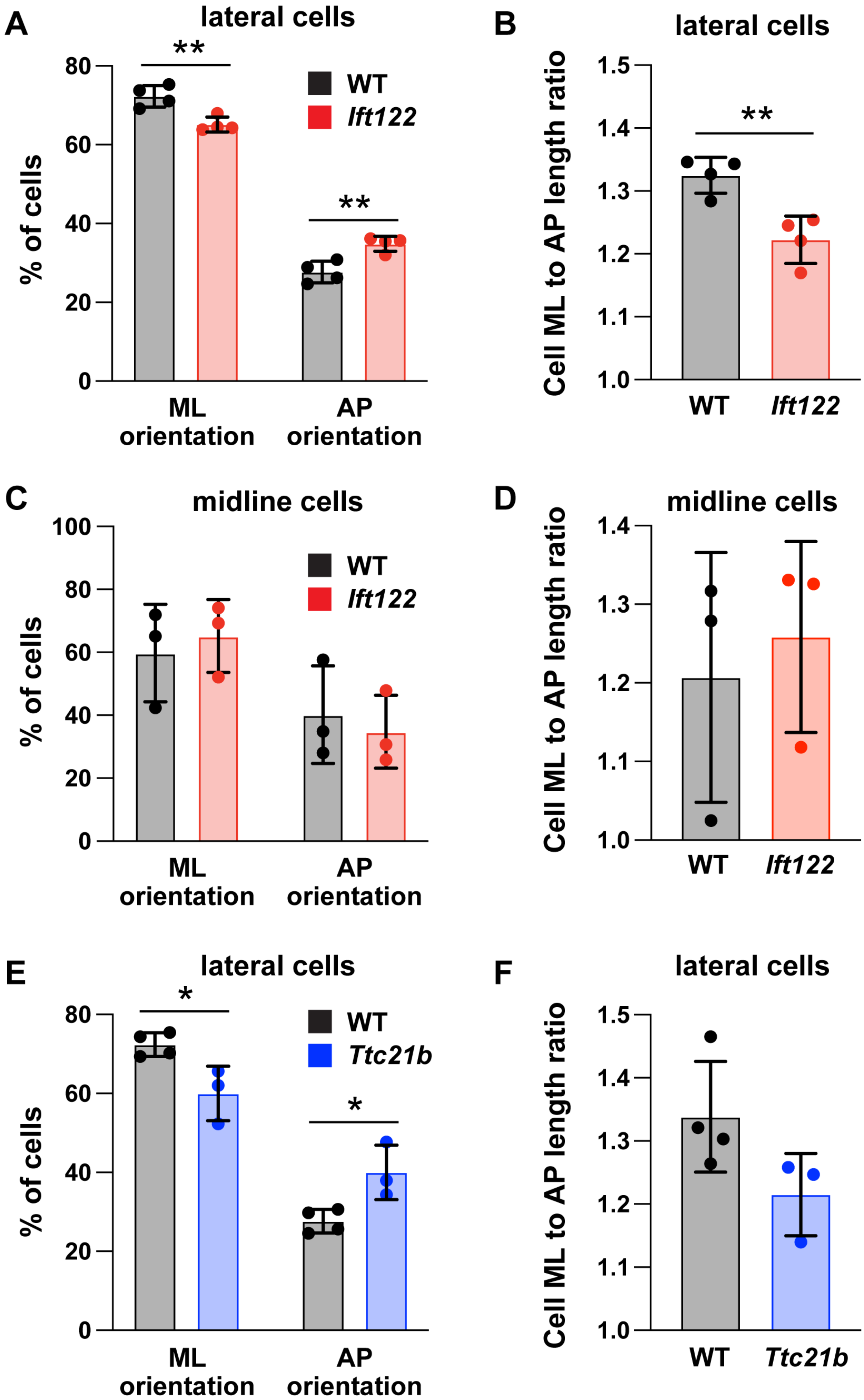

### Figure 4 - Supplement 5

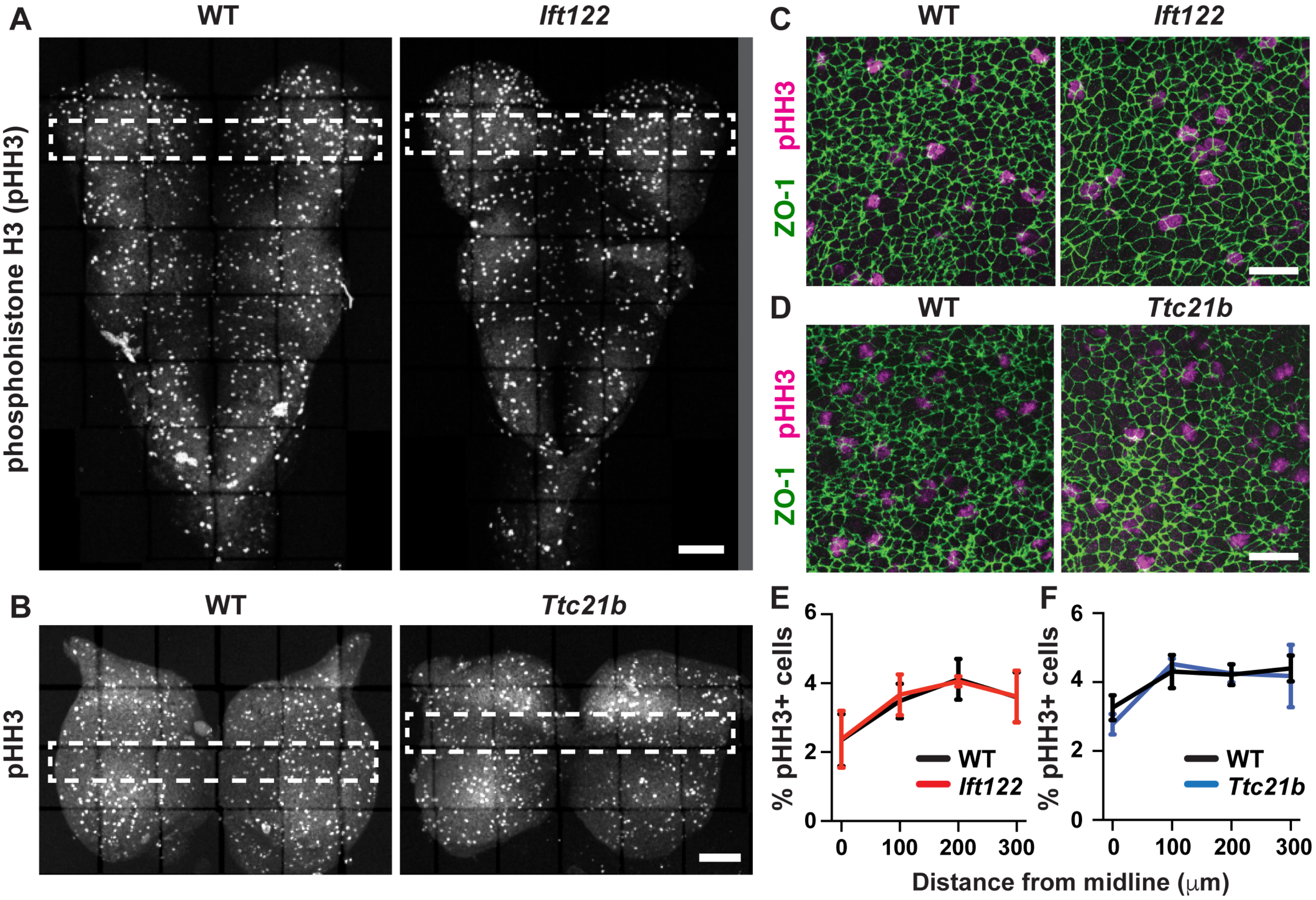

Figure 4 - Supplement 6

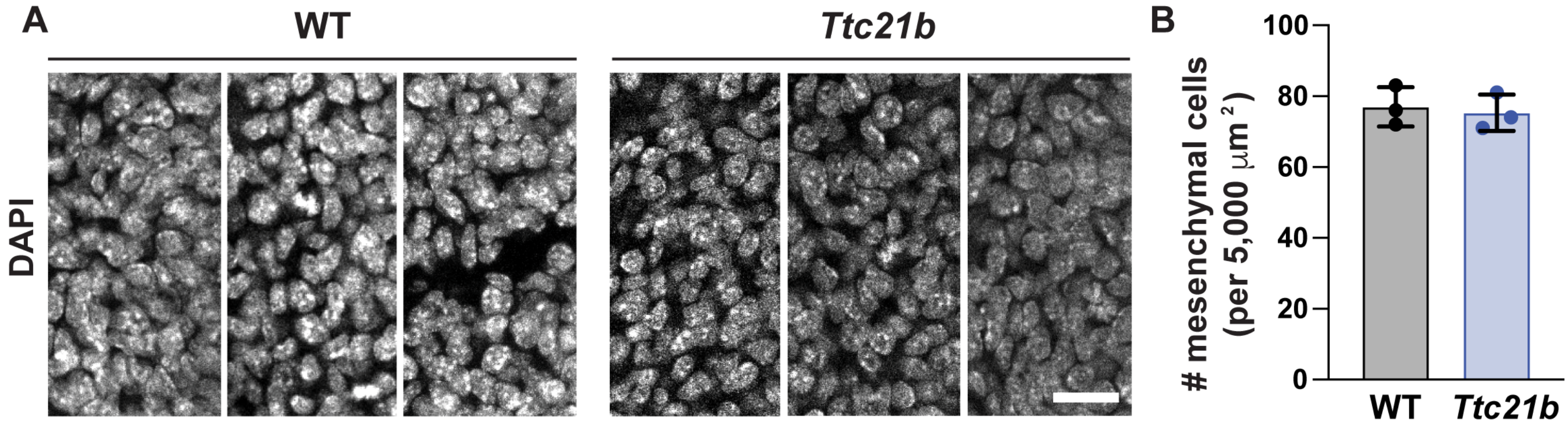

#### Figure 5 - Supplement 1

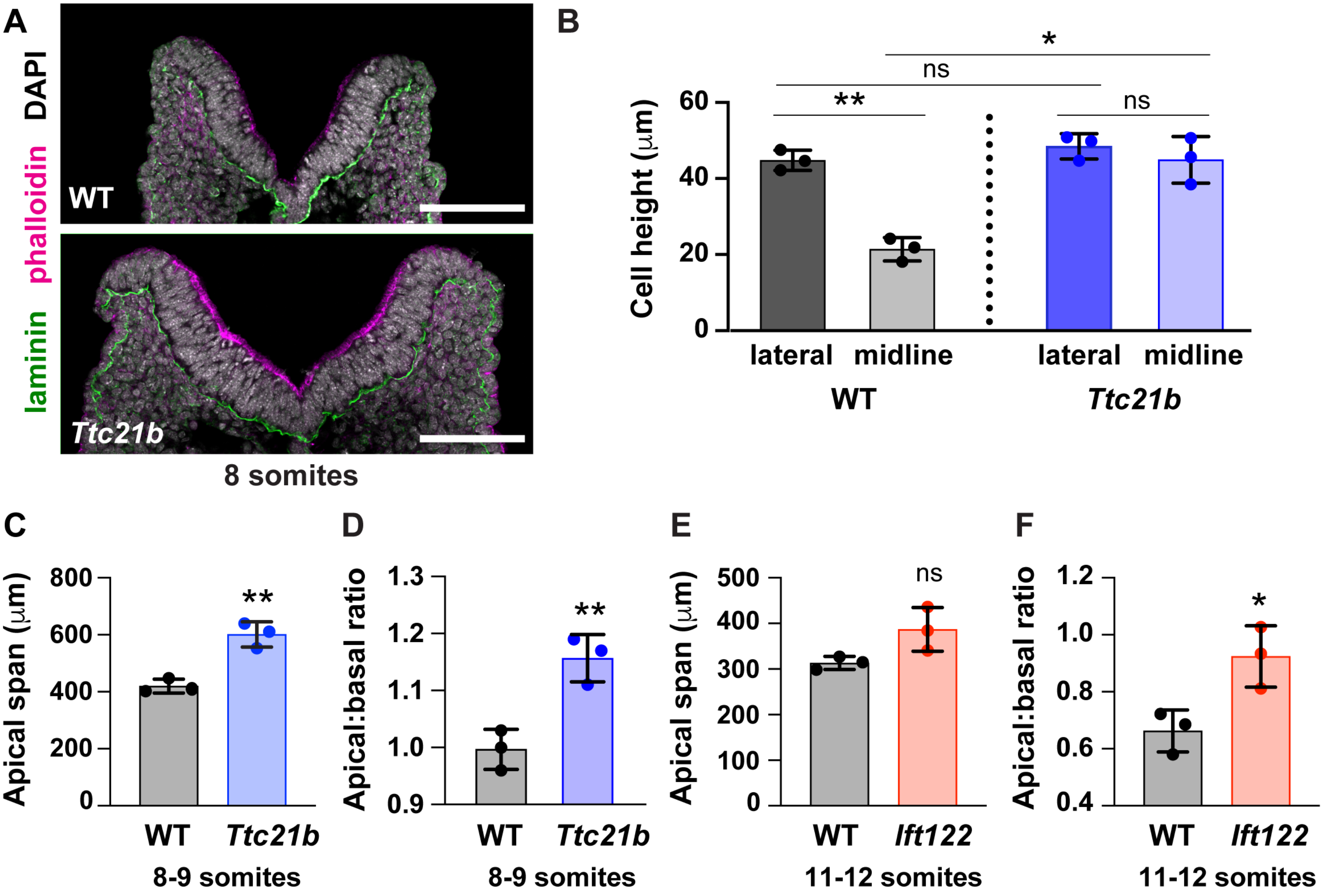

### Figure 8 - Supplement 1

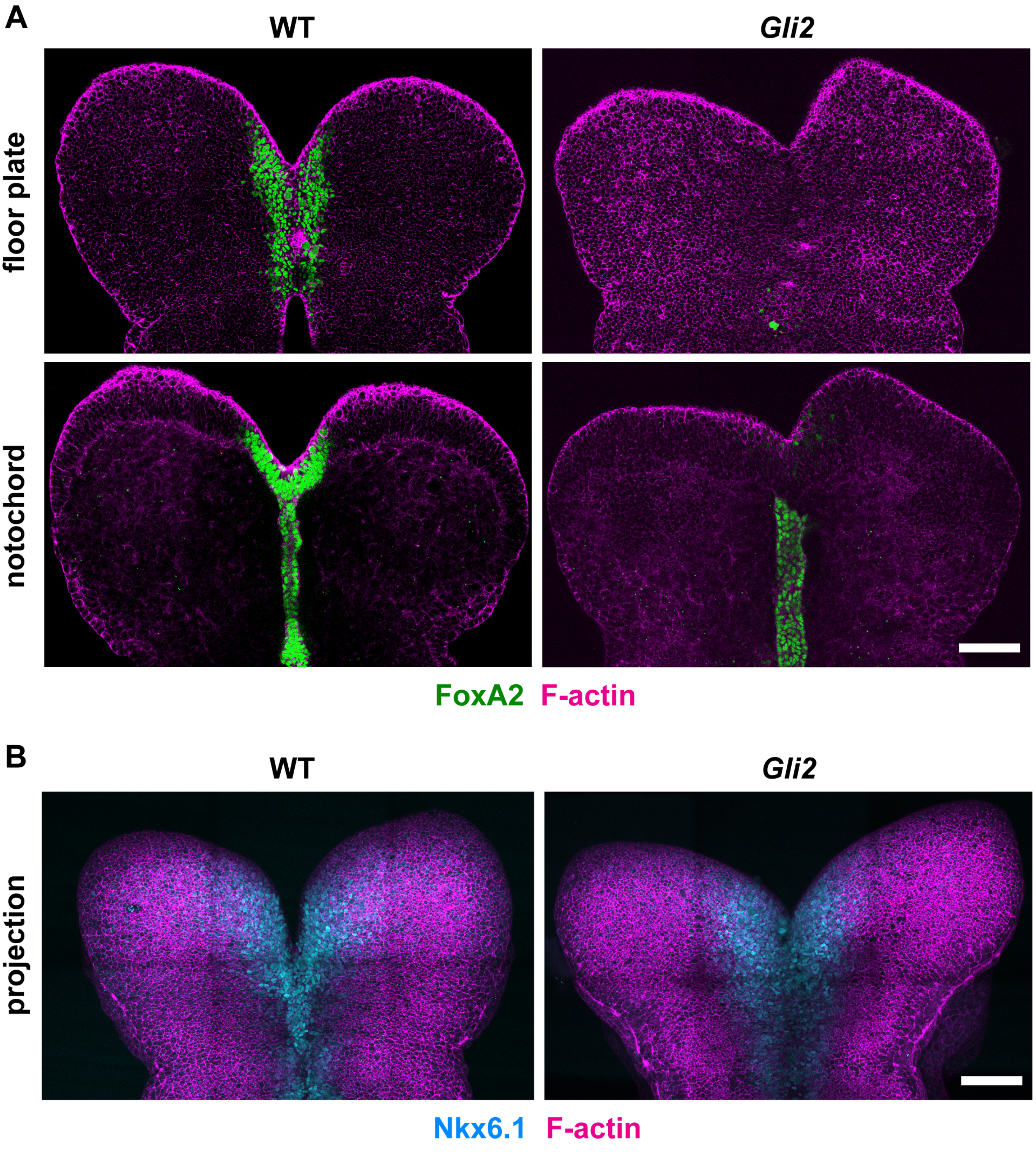

Figure 8 - Supplement 2

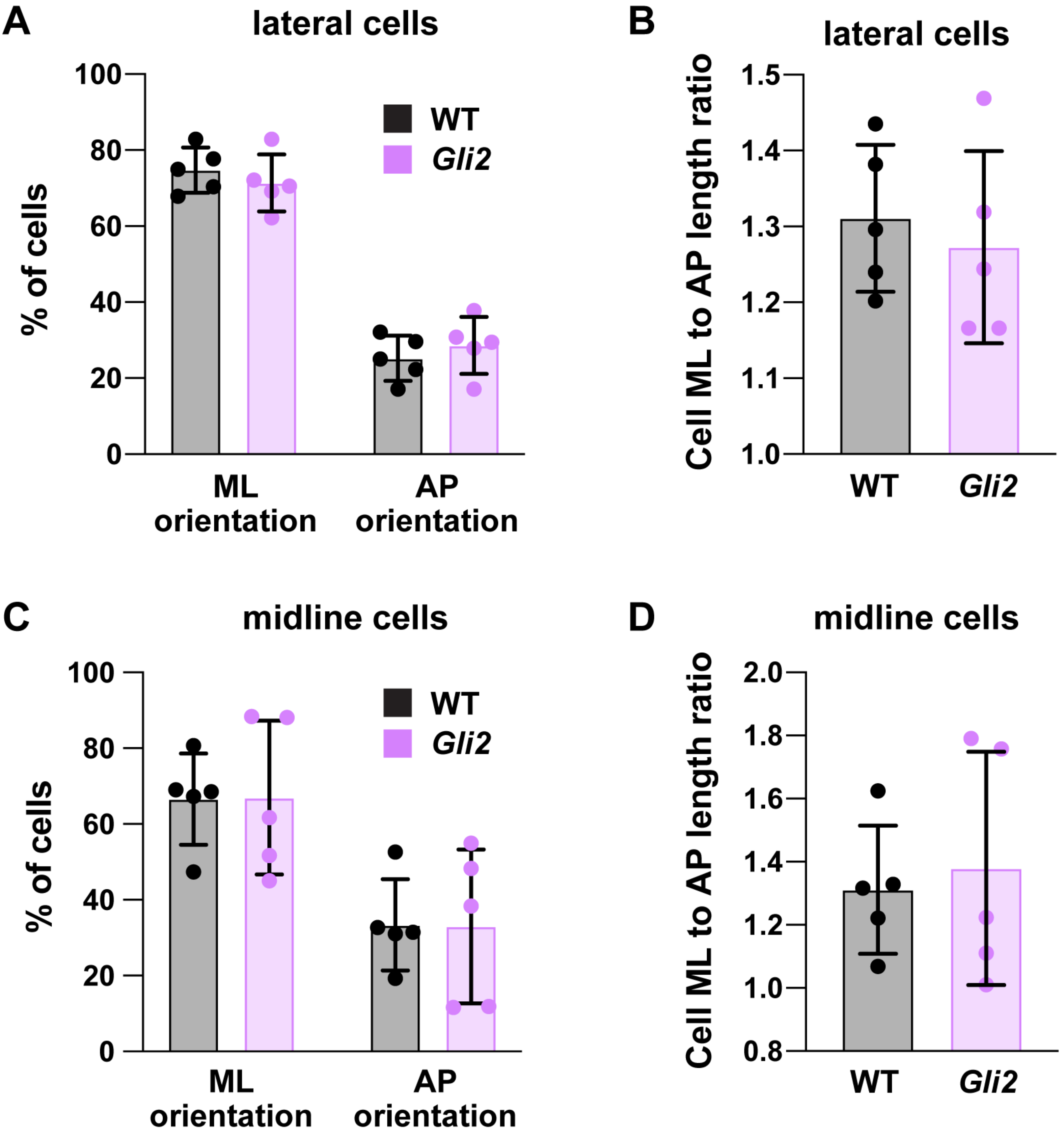

#### Figure 9 - Supplement 1

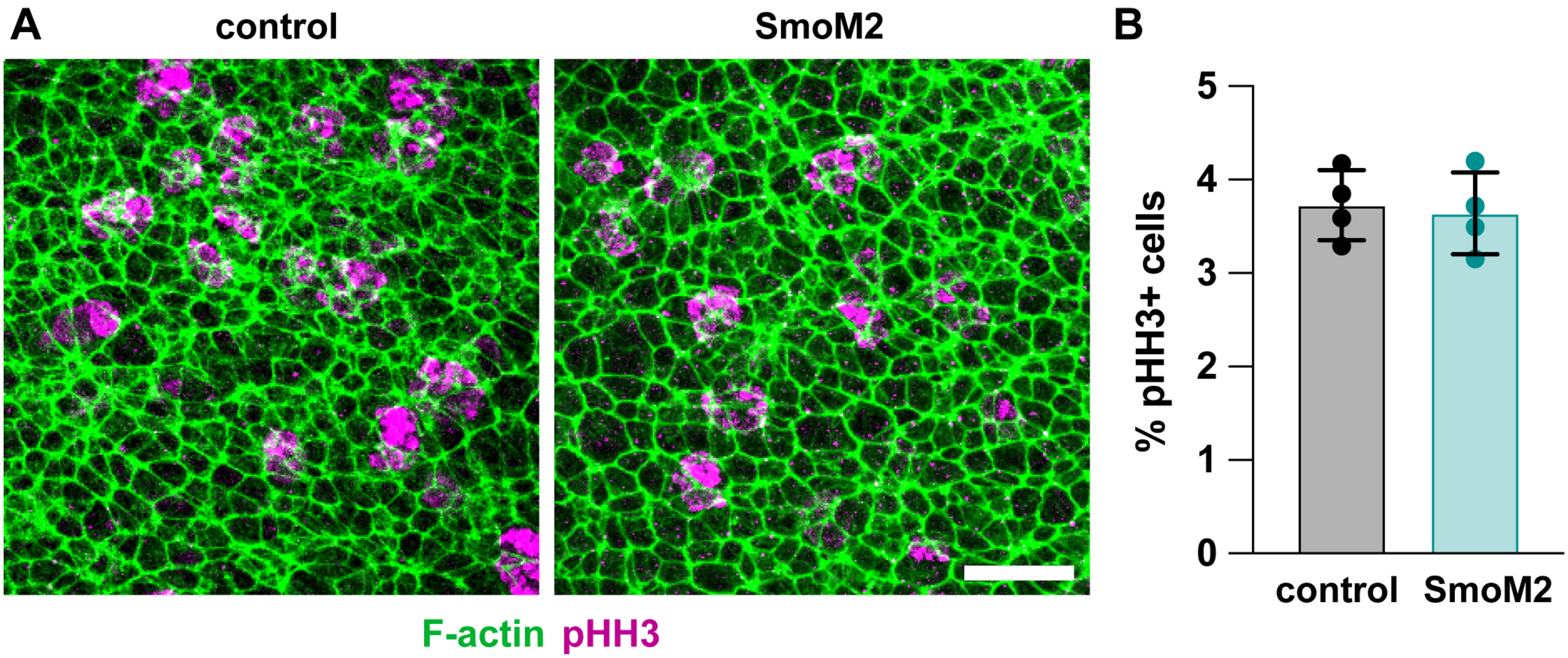

### Figure 9 - Supplement 2

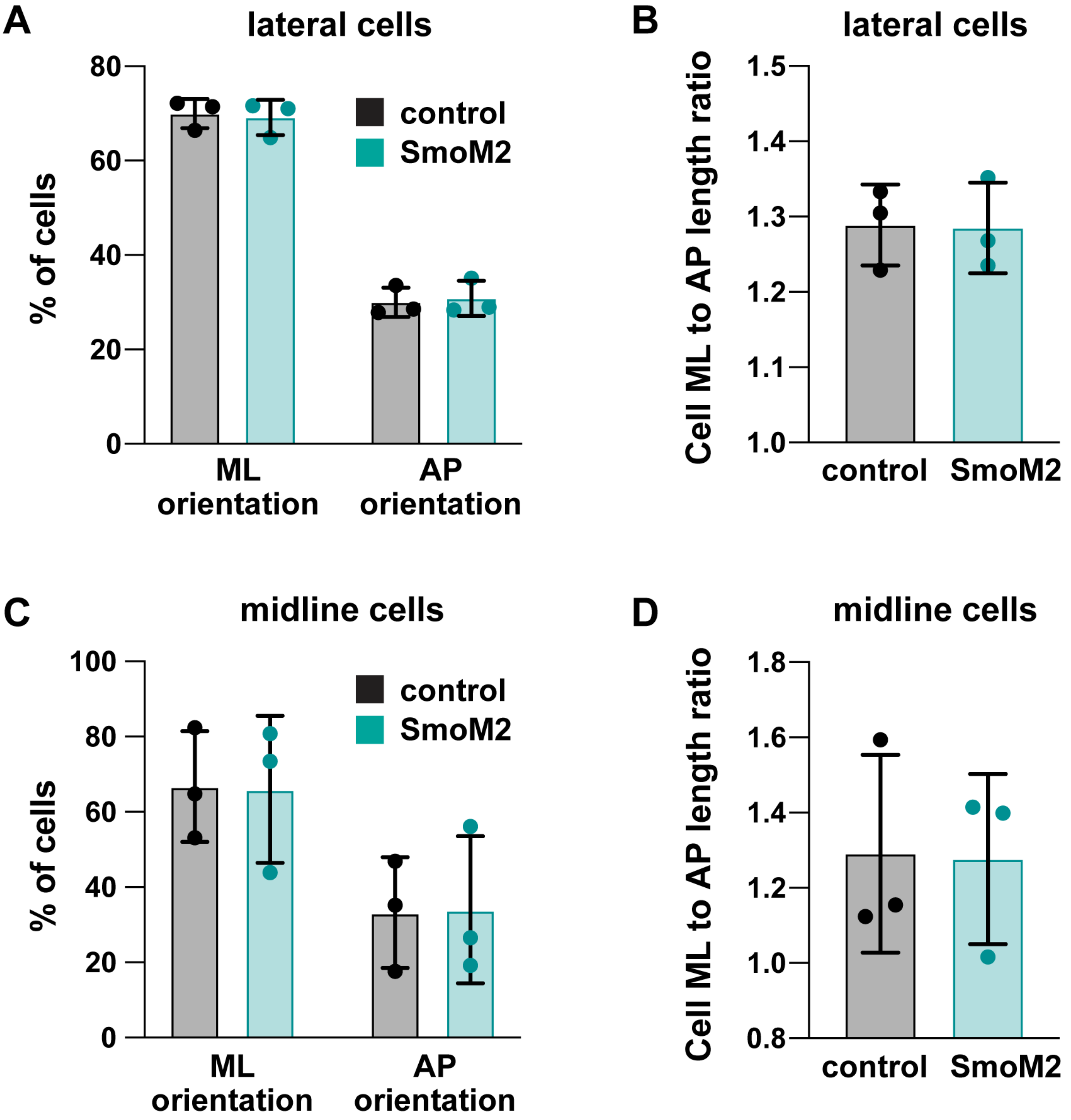
