## Supplementary File 1 for "Sonic hedgehog signaling directs patterned cell remodeling during cranial neural tube closure"

**Supplementary File 1. N values and details of statistical analyses performed.**

| Figure | Panel | n | Mean ± SD | p | Statistical test and notes |
|---|---|---|---|---|---|
| **1** | G, H | 0 somites: 3 embryos (565, 474, 488 cells)<br><br>4-6 somites: 6 embryos (857, 761, 825, 718, 667, 889 cells)<br><br>7-9 somites: 5 embryos (1000, 1270, 1198, 1047, 1377 cells) | $36.59 \pm 2.20$ µm$^2$<br><br>$24.86 \pm 2.82$ µm$^2$<br><br>$16.62 \pm 2.16$ µm$^2$ | 0 vs. 4-6: $p = 0.0027$<br><br>0 vs. 7-9: $p = 0.0006$<br><br>4-6 vs. 7-9: $p = 0.0011$ | Brown-Forsythe and Welch One-way ANOVA (Dunnett's T3 multiple comparisons)<br><br>Does not assume equal SDs |
| **2** | B, C | 0-2 somites: 3 embryos (205, 158, 174 cells)<br><br>5 somites: 3 embryos (205, 231, 197 cells)<br><br>7-9 somites: 3 embryos (192, 164, 166 cells) | $48.25 \pm 4.99$ µm$^2$<br><br>$41.22 \pm 2.22$ µm$^2$<br><br>$49.42 \pm 3.32$ µm$^2$ | 0-2 vs. 5: $p = 0.2470$<br><br>0-2 vs. 7-9: $p = 0.9781$<br><br>5 vs. 7-9: $p = 0.0878$ | Brown-Forsythe and Welch One-way ANOVA (Dunnett's T3 multiple comparisons)<br><br>Does not assume equal SDs |
| **2 S1** | B | 0 somites: 3 embryos (393, 341, 331 cells)<br><br>4-6 somites: 6 embryos (447, 595, 632, 517, 566, 484 cells)<br><br>7-9 somites: 5 embryos (552, 711, 979, 799, 915 cells) | ML: 49.81 ± 2.97%<br>AP: 50.19 ± 2.97%<br><br>ML: 69.15 ± 9.18%<br>AP: 30.85 ± 9.18%<br><br>ML: 78.17 ± 8.52%<br>AP: 21.83 ± 8.52% | 0 vs. 4-6: $p = 0.0085$<br><br>0 vs. 7-9: $p = 0.0003$<br><br>4-6 vs. 7-9: $p = 0.2228$ | Two-way ANOVA (Sidak's multiple comparisons) |
| | C | 0 somites: 3 embryos (393, 341, 331 cells) | $1.10 \pm 0.03$ | 0 vs. 4-6: $p = 0.0233$ | Brown-Forsythe and Welch One-way ANOVA |

| | | | | | |
|---|---|---|---|---|---|
| | | 4-6 somites: 6 embryos (447, 595, 632, 517, 566, 484 cells)<br><br>7-9 somites: 5 embryos (552, 711, 979, 799, 915 cells) | 1.34 ± 0.15<br><br>1.49 ± 0.19 | 0 vs. 7-9: p = 0.0322<br><br>4-6 vs. 7-9: p = 0.4490 | (Dunnett's T3 multiple comparisons)<br><br>Does not assume equal SDs |
| | E | 0 somites: 3 embryos (134, 109, 96 cells)<br><br>4-6 somites: 3 embryos (132, 148, 131 cells)<br><br>7-9 somites: 3 embryos (121, 107, 102 cells) | ML: 66.12 ± 10.41%<br>AP: 33.86 ± 10.41%<br><br>ML: 62.89 ± 16.14%<br>AP: 37.10 ± 16.14%<br><br>ML: 67.62 ± 15.89 %<br>AP: 32.37 ± 15.89% | 0 vs. 4-6: p > 0.9999<br><br>0 vs. 7-9: p > 0.9999<br><br>4-6 vs. 7-9: p > 0.9999 | Two-way ANOVA (Sidak's multiple comparisons) |
| | F | 0 somites: 3 embryos (134, 109, 96 cells)<br><br>4-6 somites: 6 embryos (132, 148, 131 cells)<br><br>7-9 somites: 5 embryos (121, 107, 102 cells) | 1.24 ± 0.16<br><br>1.28 ± 0.25<br><br>1.42 ± 0.31 | 0 vs. 5: p = 0.9939<br><br>0 vs. 7-9: p = 0.7590<br><br>5 vs. 7-9: p = 0.8903 | Brown-Forsythe and Welch One-way ANOVA (Dunnett's T3 multiple comparisons)<br><br>Does not assume equal SDs |
| **3** | B | 0-1 somite: 3 embryos<br><br>6-7 somites: 4 embryos<br><br>8-9 somites: 4 embryos | 1.34 ± 0.02<br><br>1.16 ± 0.02<br><br>0.95 ± 0.07 | 0-1 vs. 6-7: p = 0.0023<br><br>0-1 vs. 8-9: p < 0.0001<br><br>6-7 vs. 8-9: p = 0.0004 | One-way ANOVA (Tukey's multiple comparisons) |

| | | | | | |
|---|---|---|---|---|---|
| | C | 0-1 somite: 3 embryos<br><br>6-7 somites: 4 embryos<br><br>8-9 somites: 4 embryos | 32.94 ± 3.59 µm<br><br>30.84 ± 2.18 µm<br><br>51.74 ± 3.21 µm | 0-1 vs. 6-7: $p = 0.6438$<br><br>0-1 vs. 8-9: $p < 0.0001$<br><br>6-7 vs. 8-9: $p < 0.0001$ | One-way ANOVA (Tukey's multiple comparisons) |
| | D | 0-1 somite: 3 embryos<br><br>6-7 somites: 4 embryos<br><br>8-9 somites: 4 embryos | 18.38 ± 3.34 µm<br><br>18.68 ± 2.22 µm<br><br>30.40 ± 1.90 µm | 0-1 vs. 6-7: $p = 0.9858$<br><br>0-1 vs. 8-9: $p = 0.0005$<br><br>6-7 vs. 8-9: $p = 0.0004$ | One-way ANOVA (Tukey's multiple comparisons) |
| | E | 0-1 somite: 3 embryos<br><br>6-7 somites: 4 embryos<br><br>8-9 somites: 4 embryos | 1.81 ± 0.13<br><br>1.67 ± 0.25<br><br>1.71 ± 0.20 | 0-1 vs. 6-7: $p = 0.6798$<br><br>0-1 vs. 8-9: $p = 0.8234$<br><br>6-7 vs. 8-9: $p = 0.9587$ | One-way ANOVA (Tukey's multiple comparisons) |
| | G | 408 cells, 3 embryos (111, 169, 128 cells) | N/A | N/A | N/A |
| | I | 60 cells, 3 embryos (20, 20, 20 cells) | N/A | N/A | N/A |
| **3 S1** | A | 0-1 somite: 3 embryos<br><br>6-7 somites: 4 embryos<br><br>8-9 somites: 4 embryos | 652.5 ± 95.9 µm<br><br>494.9 ± 33.4 µm<br><br>406.5 ± 32.5 µm | 0-1 vs. 6-7: $p = 0.0149$<br><br>0-1 vs. 8-9: $p = 0.0011$<br><br>6-7 vs. 8-9: $p = 0.1236$ | One-way ANOVA (Tukey's multiple comparisons) |
| | B | 0-1 somite: 3 embryos<br><br>6-7 somites: 4 embryos | 486.5 ± 71.1 µm<br><br>425.6 ± 29.1 µm | 0-1 vs. 6-7: $p = 0.0976$<br><br>0-1 vs. 8-9: $p = 0.1320$ | One-way ANOVA (Tukey's multiple comparisons) |

| | | | | | |
|---|---|---|---|---|---|
| | | 8-9 somites: 4 embryos | 427.5 ± 7.5 µm | 6-7 vs. 8-9: p = 0.9998 | |
| **3 S2** | B | 3 embryos (50, 50, 50 cells) | E1: 0.17 ± 1.13<br><br>E2: 0.60 ± 1.61<br><br>E3: 0.03 ± 0.99 | N/A | N/A |
| **4** | J | WT: 4 embryos (1415, 1367, 1186, 1240 cells)<br><br>*Ift122*: 4 embryos (932, 906, 935, 757 cells) | 13.63 ± 0.82 $\mu m^2$<br><br>21.06 ± 2.00 $\mu m^2$ | p=0.0024 | Welch's t-test<br><br>Does not assume equal SDs |
| | K | WT: 5208 cells, 4 embryos<br><br>*Ift122*: 3530 cells, 4 embryos | N/A (D = 0.2763) | p<0.0001 | Kolmogorov-Smirnov |
| | L | WT: 4 embryos (1171, 1162, 1116, 1148 cells)<br><br>*Ttc21b*: 3 embryos (580, 687, 613 cells) | 15.60 ± 0.78 $\mu m^2$<br><br>30.05 ± 2.39 $\mu m^2$ | p=0.0058 | Welch's t-test<br><br>Does not assume equal SDs |
| | M | WT: 4597 cells, 4 embryos<br><br>*Ttc21b*: 1880 cells, 3 embryos | N/A (D = 0.4536) | p<0.0001 | Kolmogorov-Smirnov |
| **4 S4** | A | WT: 4 embryos (835, 1138, 1052, 990 cells)<br><br>*Ift122*: 4 embryos (715, 867, 626, 563 cells) | WT ML: 72.32 ± 2.74%<br>WT AP: 27.68 ± 2.74%<br><br>*Ift122* ML: 65.14 ± 1.89%<br>*Ift122* AP: 34.86 ± 1.89% | WT vs. *Ift122* ML: p = 0.0020<br><br>WT vs. *Ift122* AP: p = 0.0020 | Two-way ANOVA (Sidak's multiple comparisons) |
| | B | WT: 4 embryos (835, 1138, 1052, 990 cells) | 1.33 ± 0.03<br><br>1.22 ± 0.04 | p = 0.0058 | Welch's t-test<br><br>Does not |

| | | | | | |
|---|---|---|---|---|---|
| | | *Ift122*: 4 embryos (715, 867, 626, 563 cells) | | | assume equal SDs |
| | C | WT: 3 embryos (146, 170, 195 cells)<br><br>*Ift122*: 3 embryos (372, 599, 414 cells) | WT ML: 59.80 ± 15.49%<br>WT AP: 40.20 ± 15.49%<br><br>*Ift122* ML: 65.18 ± 11.60%<br>*Ift122* AP: 34.82 ± 11.60% | WT vs. *Ift122* ML: p = 0.8725<br><br>WT vs. *Ift122* AP: p = 0.8725 | Two-way ANOVA (Sidak's multiple comparisons) |
| | D | WT: 3 embryos (146, 170, 195 cells)<br><br>*Ift122*: 3 embryos (372, 599, 414 cells) | 1.20 ± 0.16<br><br>1.26 ± 0.12 | p = 0.6810 | Welch's t-test<br><br>Does not assume equal SDs |
| | E | WT: 4 embryos (913, 898, 870, 912 cells)<br><br>*Ttc21b*: 3 embryos (415, 489, 411 cells) | WT ML: 72.36 ± 3.00%<br>WT AP: 27.64 ± 3.00%<br><br>*Ttc21b* ML: 59.99 ± 6.91%<br>*Ttc21b* AP: 40.01 ± 6.91% | WT vs. *Ttc21b* ML: p = 0.0167<br><br>WT vs. *Ttc21b* AP: p = 0.0167 | Two-way ANOVA (Sidak's multiple comparisons) |
| | F | WT: 4 embryos (913, 898, 870, 912 cells)<br><br>*Ttc21b*: 3 embryos (415, 489, 411 cells) | 1.34 ± 0.09<br><br>1.22 ± 0.07 | p = 0.0863 | Welch's t-test<br><br>Does not assume equal SDs |
| **4 S5** | E | WT:<br>3 embryos | WT 0-100:<br>2.34 ± 0.76 %<br><br>WT 100-200:<br>3.48 ± 0.50 %<br><br>WT 200-300: 4.12 ± 0.59 %<br><br>WT 300-400:<br>3.60 ± 0.72 % | WT vs. *Ift122*<br><br>0-100:<br>p > 0.9999<br><br>100-200:<br>p = 0.9950<br><br>200-300:<br>p > 0.9999 | Two-way ANOVA (Sidak's multiple comparisons) |

| | | | | | |
|---|---|---|---|---|---|
| | | *Ift122*:<br>3 embryos | *Ift122* 0-100:<br>2.37 ± 0.83 %<br><br>*Ift122* 100-200:<br>3.67 ± 0.59 %<br><br>*Ift122* 200-300:<br>4.05 ± 0.15 %<br><br>*Ift122* 300-400:<br>3.62 ± 0.75 % | 300-400:<br>p > 0.9999 | |
| | F | WT:<br>3 embryos<br><br>*Ttc21b*:<br>3 embryos | WT 0-100:<br>3.25 ± 0.36 %<br><br>WT 100-200:<br>4.31 ± 0.48 %<br><br>WT 200-300: 4.22 ± 0.31 %<br><br>WT 300-400:<br>4.40 ± 0.38 %<br><br>*Ttc21b* 0-100:<br>2.78 ± 0.29 %<br><br>*Ttc21b* 100-200:<br>4.52 ± 0.16 %<br><br>*Ttc21b* 200-300:<br>4.25 ± 0.26 %<br><br>*Ttc21b* 300-400:<br>4.18 ± 0.90 % | WT vs. *Ttc21b*<br><br>0-100:<br>p = 0.6082<br><br>100-200:<br>p = 0.9633<br><br>200-300:<br>p > 0.9999<br><br>300-400:<br>p > 0.9618 | Two-way ANOVA (Sidak's multiple comparisons) |
| **4 S6** | B | WT: 3 embryos<br><br>*Ttc21b*: 3 embryos | 77.00 ± 5.57 cells<br><br>75.33 ± 5.13 cells | p = 0.7225 | Welch's t-test<br><br>Does not assume equal SDs |
| **5** | C, D | WT lateral: 4 embryos (data from Figure 4K)<br><br>WT midline: 3 | N/A | N/A | N/A |

| | | | | | |
|---|---|---|---|---|---|
| | | embryos (276, 316, 285 cells)<br><br>*Ift122* lateral: 4 embryos (data from Figure 4K)<br><br>*Ift122* midline: 3 embryos (516, 441, 539 cells) | | | |
| | E | WT lateral: 4 embryos (data from Figure 4J)<br><br>WT midline: 3 embryos (276, 316, 285 cells)<br><br>*Ift122* lateral: 4 embryos (data from Figure 4J)<br><br>*Ift122* midline: 3 embryos (516, 441, 539 cells) | WT:<br>lateral: 13.63 ± 0.82 $\mu m^2$<br>midline: 32.45 ± 2.18 $\mu m^2$<br><br>*Ift122*:<br>lateral: 21.06 ± 2.00 $\mu m^2$<br>midline: 19.23 ± 2.37 $\mu m^2$ | WT lat vs. WT mid: $p = 0.0149$<br><br>*Ift122* lat vs. *Ift122* mid: $p = 0.8383$<br><br>WT lat vs. *Ift122* lat: $p = 0.0100$<br><br>WT lat vs. *Ift122* mid: $p = 0.1723$<br><br>WT mid vs. *Ift122* lat: $p = 0.0091$<br>WT mid vs. *Ift122* mid: $p = 0.0090$ | Brown-Forsythe and Welch One-way ANOVA (Dunnett's T3 multiple comparisons)<br><br>Does not assume equal SDs |
| | G | WT: 3 embryos<br>*Ift122*: 3 embryos | N/A | N/A | N/A |
| | H | WT: 3 embryos<br>*Ttc21b*: 3 embryos | N/A | N/A | N/A |
| | I | WT (lateral and midline): 3 embryos | WT:<br>lateral: 59.80 ± 1.90 μm<br>midline: 26.63 ± 2.97 μm | WT lat vs. WT mid: $p = 0.0019$<br><br>*Ift122* lat vs. *Ift122* mid: $p = 0.4142$ | Brown-Forsythe and Welch One-way ANOVA (Dunnett's T3 multiple comparisons) |

| | | | | | |
|---|---|---|---|---|---|
| | | *Ift122* (lateral and midline): 3 embryos | *Ift122*: lateral: 51.13 ± 1.10 µm midline: 45.81 ± 3.96 µm | WT lat vs. *Ift122* lat: p = 0.0239<br><br>WT lat vs. *Ift122* mid: p = 0.0433<br><br>WT mid vs. *Ift122* lat: p = 0.0034<br><br>WT mid vs. *Ift122* mid: p = 0.0111 | Does not assume equal SDs |
| **5 S1** | B | WT: 3 embryos<br><br>*Ttc21b*: 3 embryos | WT lateral: 44.77 ± 2.66 µm<br><br>WT midline: 21.40 ± 3.06 µm<br><br>*Ttc21b* lateral: 48.44 ± 3.32 µm<br><br>*Ttc21b* midline: 44.91 ± 6.12 µm | WT lat vs. WT mid: p = 0.0025<br><br>WT lat vs *Ttc21b* lat: p = 0.6285<br><br>WT lat vs. *Ttc21b* mid: p > 0.9999<br><br>WT mid vs. *Ttc21b* lat: p = 0.0021<br><br>WT mid vs. *Ttc21b* mid: p = 0.0352<br><br>*Ttc21b* lat vs. *Ttc21b* mid: p = 0.9140 | Brown-Forsythe and Welch One-way ANOVA (Dunnett's T3 multiple comparisons)<br><br>Does not assume equal SDs |
| | C | WT: 3 embryos<br><br>*Ttc21b*: 3 embryos | 420.0 ± 24.2 µm<br><br>601.3 ± 44.1 µm | p = 0.0075 | Welch's t-test<br><br>Does not assume equal SDs |
| | D | WT: 3 embryos | 1.00 ± 0.04 | p = 0.0076 | Welch's t-test |

| | | | | | |
|---|---|---|---|---|---|
| | | *Ttc21b*: 3 embryos | 1.16 ± 0.04 | | Does not assume equal SDs |
| | E | WT: 3 embryos<br><br>*Ift122*: 3 embryos | 313.3 ± 14.2 µm<br><br>387.0 ± 47.9 µm | $p = 0.1065$ | Welch's t-test<br><br>Does not assume equal SDs |
| | F | WT: 3 embryos<br><br>*Ift122:* 3 embryos | 0.66 ± 0.07<br><br>0.92 ± 0.11 | p = 0. 0314 | Welch's t-test<br><br>Does not assume equal SDs |
| **6** | B (*Ift122*) | WT:<br>86 cables, 3 embryos<br><br>*Ift122*:<br>36 cables, 3 embryos | WT circular mean: 24.2°<br><br>*Ift122* circular mean: 41.5° | $P < 0.05$ | Watson nonparametric two-sample test for homogeneity |
| | B (*Ttc21b*) | WT:<br>84 cables, 3 embryos<br><br>*Ttc21b:*<br>29 cables, 3 embryos | WT circular mean: 26.4°<br><br>*Ttc21b* circular mean: 44.3° | $0.05 < P < 0.10$ | Watson nonparametric two-sample test for homogeneity |
| | C (*Ift122)* | WT:<br>3 embryos<br><br>*Ift122*:<br>3 embryos | 28.67 ± 6.11 cables<br><br>12.33 ± 3.06 cables | $p = 0.0266$ | Welch's t-test<br><br>Does not assume equal SDs |
| | C (*Ttc21b*) | WT:<br>3 embryos<br><br>*Ttc21b*:<br>3 embryos | 28.0 ± 1.7 cables<br><br>9.7 ± 5.0 cables | $p = 0.0160$ | Welch's t-test<br><br>Does not assume equal SDs |
| | D (*Ift122*) | WT:<br>3 embryos<br><br>*Ift122*:<br>3 embryos | 50.7 ± 9.3 cables<br><br>33.3 ± 4.0 cables | $p = 0.06668$ | Welch's t-test<br><br>Does not assume equal SDs |

| | | | | | |
|---|---|---|---|---|---|
| | D (*Ttc21b*) | WT:<br>3 embryos<br><br>*Ttc21b*:<br>3 embryos | 43.4 ± 3.8 cables<br><br><br>33.7 ± 4.2 cables | $p = 0.0414$ | Welch's t-test<br><br>Does not assume equal SDs |
| | F (*Ift122*) | WT:<br>151 cables, 3 embryos<br><br>*Ift122*:<br>100 cables, 3 embryos | WT circular mean: 34.7°<br><br>*Ift122* circular mean: 45.9° | $P < 0.01$ | Watson nonparametric two-sample test for homogeneity |
| | F (*Ttc21b*) | WT:<br>130 cables, 3 embryos<br><br>*Ttc21b:*<br>101 cables, 3 embryos | WT circular mean: 28.8°<br><br>*Ttc21b* circular mean: 40.8° | $P < 0.001$ | Watson nonparametric two-sample test for homogeneity |
| | G (*Ift122*) | WT: 3 embryos<br>(50, 50, 50 cells)<br><br>*Ift122*: 3 embryos<br>(50, 50, 50 cells) | N/A | N/A | N/A |
| | H (*Ttc21b*) | WT: 3 embryos<br>(50, 50, 50 cells)<br><br>*Ttc21b*: 3 embryos<br>(50, 50, 50 cells) | N/A | N/A | N/A |
| **7** | B | WT: 3 embryos<br><br>*Ift122*: 3 embryos | N/A | N/A | N/A |
| | C | WT: 3 embryos<br><br>*Ttc21b*: 3 embryos | N/A | N/A | N/A |
| **8** | B, C | WT: 5 embryos<br>(1129, 1011, 1047, 1269, 1105 cells)<br><br>*Gli2*: 5 embryos<br>(1075, 967, 1067, 1017, 1097 cells) | 17.39 ± 0.78 $\mu m^2$<br><br><br>16.96 ± 1.57 $\mu m^2$ | $p = 0.6069$ | Welch's t-test<br><br>Does not assume equal SDs |

| | | | | | |
|---|---|---|---|---|---|
| | E, F | WT: 5 embryos (233, 220, 161, 342, 310 cells)<br><br>*Gli2*: 5 embryos (228, 257, 277, 171, 174 cells) | 47.66 ± 7.20 $\mu m^2$<br><br>32.24 ± 5.98 $\mu m^2$ | p = 0.0066 | Welch's t-test<br><br>Does not assume equal SDs |
| | H | WT: 5 embryos<br><br>*Gli2*: 5 embryos | WT lateral: 59.54 ± 2.14 µm<br><br>WT midline: 29.90 ± 1.80 µm<br><br>*Gli2* lateral: 57.37 ± 5.95 µm<br><br>*Gli2* midline: 50.01 ± 3.04 µm | WT lat vs. WT mid: p < 0.0001<br><br>WT lat vs. *Gli2* lat: p = 0.9555<br><br>WT lat vs. *Gli2* mid: p = 0.0038<br><br>WT mid vs. *Gli2* lat: p = 0.0009<br><br>WT mid vs. *Gli2* mid: p < 0.0001<br><br>*Gli2* lat vs. *Gli2* mid: p = 0.2100 | Brown-Forsythe and Welch One-way ANOVA (Dunnett's T3 multiple comparisons)<br><br>Does not assume equal SDs |
| **8S2** | A | WT: 5 embryos (903, 796, 823, 792, 860 cells)<br><br>*Gli2*: 5 embryos (1023, 867, 831, 738, 836 cells) | WT ML: 74.76 ± 5.95%<br>WT AP: 25.24 ± 5.95%<br><br>*Gli2* ML: 71.73 ± 7.47%<br>*Gli2* AP: 28.63 ± 7.47% | WT vs. *Gli2* ML: p = 0.6853<br><br>WT vs. *Gli2* AP: p = 0.6853 | Two-way ANOVA (Sidak's multiple comparisons) |
| | B | WT: 5 embryos (903, 796, 823, 792, 860 cells)<br><br>*Gli2*: 5 embryos (1023, 867, 831, 738, 836 cells) | 1.31 ± 0.10<br><br>1.27 ± 0.13 | p = 0.6079 | Welch's t-test<br><br>Does not assume equal SDs |
| | C | WT: 5 embryos (260, 173, 88, 105, 100 cells) | WT ML: 66.59 ± 12.01% | WT vs. *Gli2* ML: p = 0.9991 | Two-way ANOVA (Sidak's |

| | | | | | |
|---|---|---|---|---|---|
| | | *Gli2*: 5 embryos (244, 325, 180, 177, 146 cells) | WT AP: 33.41 ± 12.01%<br><br>*Gli2* ML: 66.99 ± 20.28%<br>*Gli2* AP: 33.01 ± 20.28% | WT vs. *Gli2* AP: p = 0.9991 | multiple comparisons) |
| | D | WT: 5 embryos (260, 173, 88, 105, 100 cells)<br><br>*Gli2*: 5 embryos (244, 325, 180, 177, 146 cells) | 1.31 ± 0.20<br><br>1.38 ± 0.37 | p = 0.7324 | Welch's t-test<br><br>Does not assume equal SDs |
| **9** | H | control: 3 embryos (1140, 1104, 1395 cells)<br><br>Wnt1-Cre2 > SmoM2: 3 embryos (763, 867, 723 cells) | 15.15 ± 1.61 $\mu m^2$<br><br>22.69 ± 2.07 $\mu m^2$ | p = 0.0089 | Welch's t-test<br><br>Does not assume equal SDs |
| | I | control: 3639 cells, 3 embryos<br><br>Wnt1-Cre2 > SmoM2: 2353 cells, 3 embryos | N/A (D = 0.3002) | p < 0.0001 | Kolmogorov-Smirnov |
| | J | control: 3 embryos (225, 183, 189 cells)<br><br>Wnt1-Cre2 > SmoM2: 3 embryos (188, 176, 224 cells) | 43.76 ± 4.85 $\mu m^2$<br><br>43.56 ± 3.81 $\mu m^2$ | p = 0.9556 | Welch's t-test<br><br>Does not assume equal SDs |
| | K | control: 597 cells, 3 embryos<br><br>Wnt1-Cre2 > SmoM2: 588 cells, 3 embryos | N/A (D = 0.0320) | p = 0.9215 | Kolmogorov-Smirnov |
| **9 S1** | B | control: 4 embryos<br><br>Wnt1-Cre2 > SmoM2: 4 embryos | 3.73 ± 0.38 %<br><br>3.64 ± 0.44 % | p = 0.7798 | Welch's t-test<br><br>Does not assume equal SDs |

| | | | | | |
|---|---|---|---|---|---|
| **9S2** | A | control: 3 embryos (887, 912, 1132 cells)<br><br>SmoM2: 3 embryos (567, 655, 753 cells) | control ML: 69.99 ± 3.13%<br>control AP: 30.01 ± 3.13%<br><br>SmoM2 ML: 69.19 ± 3.72%<br>SmoM2 AP: 30.81 ± 3.72% | control vs. SmoM2 ML: p = 0.9519<br><br>control vs. SmoM2 AP: p = 0.9519 | Two-way ANOVA (Sidak's multiple comparisons) |
| | B | control: 3 embryos (887, 912, 1132 cells)<br><br>SmoM2: 3 embryos (567, 655, 753 cells) | 1.29 ± 0.05<br><br>1.29 ± 0.60 | p = 0.9359 | Welch's t-test<br><br>Does not assume equal SDs |
| | C | control: 3 embryos (85, 143, 125 cells)<br><br>SmoM2: 3 embryos (146, 78, 144 cells) | control ML: 66.77 ± 14.70%<br>control AP: 33.23 ± 14.70%<br><br>SmoM2 ML: 66.01 ± 19.55%<br>SmoM2 AP: 33.99 ± 19.55% | control vs SmoM2 ML: p = 0.9983<br><br>control vs SmoM2 AP: p = 0.9983 | Two-way ANOVA (Sidak's multiple comparisons) |
| | D | control: 3 embryos (85, 143, 125 cells)<br><br>SmoM2: 3 embryos (146, 78, 144 cells) | 1.31 ± 0.20<br><br>1.38 ± 0.37 | p = 0.7324 | Welch's t-test<br><br>Does not assume equal SDs |
